# Structure-function analysis reveals a DNA polymerization-independent role for mitochondrial DNA polymerase IC in African trypanosomes

**DOI:** 10.1101/730523

**Authors:** Jonathan C Miller, Stephanie B Delzell, Jeniffer Concepción-Acevedo, Michael J Boucher, Michele M Klingbeil

## Abstract

The mitochondrial DNA of *Trypanosoma brucei* and related parasites is a catenated network containing thousands of minicircles and tens of maxicircles called kinetoplast DNA (kDNA). Replication of the single nucleoid requires at least three DNA polymerases (POLIB, POLIC, and POLID) each having discrete localization near the kDNA during S phase. POLIB and POLID have roles in minicircle replication while the specific role of POLIC in kDNA maintenance is less clear. Here, we use an RNAi-complementation system to dissect the functions of the distinct POLIC domains: the conserved family A DNA polymerase domain (POLA) and the uncharacterized N-terminal region (UCR). While RNAi complementation with wild-type POLIC restored kDNA content and cell cycle localization, active site point mutations in the POLA domain impaired minicircle replication similarly to POLIB and POLID depletions. Complementation with the POLA domain alone abolished POLIC foci formation and partially rescued the RNAi phenotype. Furthermore, we provide evidence of a crucial role for the UCR in cell cycle localization and segregation of kDNA daughter networks. This is the first report of a DNA polymerase that impacts DNA segregation.

**Summary statement:** Mitochondrial DNA segregation in African trypanosomes is supported by a dual-functioning DNA polymerase.

## INTRODUCTION

Maintenance of mitochondrial DNA (mtDNA) is vital for most eukaryotic cells. Accumulation of mutations or failure to maintain copy number can result in loss of mtDNA-encoded proteins essential for energy generation and mitochondrial dysfunction that is associated with aging, metabolic disease, neurodegenerative disorders, and cancer (Viscomi and Zeviani, 2017). As a result, mtDNA maintenance is a focal point of biomedical research. Despite decades of investigation, the mechanisms that regulate replication, copy number maintenance and segregation of mtDNA remain unknown.

Although many studies on mitochondrial biology have focused on a few eukaryotic lineages (yeast and mammals), it is now clear that great diversity exists in mitochondrial genomes and proteomes across eukaryotes. (Chen and Butow, 2005; Gray, 2015; Zíková et al., 2016). Eukaryotic microbes constitute much of this evolutionary diversity, and the parasitic protists of medical importance are by far the most well-studied. One of the most intriguing and structurally complex mitochondrial genomes is found in trypanosomatid protists such as *Trypanosoma brucei*, the parasite responsible for African sleeping sickness. Trypanosomatids harbor a single mitochondrion (often branched) and the mtDNA consists of two genetic elements, the minicircles and maxicircles that are catenated into a network called kinetoplast DNA (kDNA) (Jensen and Englund, 2012; Verner et al., 2015). *In vivo*, kDNA is condensed into a single disk-shaped nucleoid always present near the flagellar basal body and is replicated once every cell cycle in near synchrony with other single unit organelles, including the nucleus and flagellum (Woodward and Gull, 1990). The topological complexity of kDNA provides a striking reporter system in which lesions in the network and disruption of replication intermediates help to define mechanistic defects and therefore represents a good model to study mtDNA dynamics.

Variations on kDNA structure and the sizes of minicircles and maxicircles exist (for review see Lukes et al., 2002). In *T. brucei*, the 23 kb maxicircles (approximately 25 copies) are considered homologs to mammalian mtDNA as they contain similar protein-coding genes for respiratory complex subunits and rRNAs. However, a majority of the maxicircle genes are cryptic and undergo an extensive post-transcriptional RNA editing process of uridine insertion and deletion to generate translatable mRNAs. This process is mediated by minicircle-encoded gRNAs (for reviews see (Read et al., 2016; Stuart et al., 2005). Therefore, both genetic elements are essential for fitness and completion of the parasite’s digenetic life cycle (Dewar et al., 2018).

Moreover, the ∼5000 minicircles (1 kb) constitute the majority of the network with each minicircle Hopf-linked to three others (Chen et al., 1995). The maxicircles are linked to the minicircles and are also likely interlocked with other maxicircles forming a network within a network (Shapiro, 1993). How this complex structure is replicated and segregated are areas of focused study. (for reviews see (Jensen and Englund, 2012; Povelones, 2014; Schneider and Ochsenreiter, 2018).

One key feature is the topoisomerase II-mediated release of minicircle monomers to replicate free of the network. Briefly, unreplicated covalently closed (CC) minicircles are released into a region between the kDNA disk and mitochondrial membrane nearest the flagellar basal body, the kinetoflagellar zone (KFZ), where initiation and synthesis occur via unidirectional theta replication (Abu-Elneel et al., 2001; Melendy et al., 1988). Later stages such as Okazaki fragment processing and topoisomerase-mediated attachment of minicircle progeny occur in two protein assemblies located at opposite poles of the network periphery called antipodal sites (AS) (Hines et al., 2001; Melendy et al., 1988). Minicircle progeny are attached to the network while still containing at least one nick or gap (N/G); a presumed counting mechanism that distinguishes between replicated versus non-replicated minicircles (Ntambi et al., 1986). The process results in a spatial and temporal separation of replication events. Maxicircles replicate via theta replication within the network and progeny also contain N/G (Carpenter and Englund, 1995; Liu et al., 2009). During the later stages of network replication the remaining nicks and gaps are repaired and the network undergoes topological remodeling prior to segregation.

A second key feature is the tripartite attachment complex (TAC) that physically links the kDNA network to the basal body of the flagellum and mediates segregation of the progeny kDNA networks (Ogbadoyi, Robinson, Gull, 2003). The TAC is composed of three distinct regions: the exclusion zone filaments (EZF) that extend from the basal body to the mitochondrial outer membrane, the differentiated mitochondrial membrane (DM) located between the basal body and the kDNA disk, and the unilateral filaments (ULF) that extend from the mitochondrial inner membrane to the kDNA network (in the KFZ). Elegant electron microscopy (EM) cytochemistry studies defined inner and outer ULF domains (Gluenz et al., 2007) that could represent subdomains; one that occupies replication processes and others that occupy structural or segregation processes. Lastly, several components of each region have been identified and genetic studies indicate a hierarchical organization for TAC biogenesis starting with components that are most proximal to the basal body (Hoffmann et al., 2018; Schneider and Ochsenreiter, 2018). One core TAC component, TAC102, localizes to the ULF and is essential for proper kDNA segregation, although it does not directly interact with the kDNA network it is the most proximal ULF component (Jakob et al., 2016; Trikin et al., 2016).

A third feature is the multiplicity of proteins with similar activities but non-redundant roles in kDNA maintenance. Among the many additional factors not present in mammalian nucleoids are 2 topoisomerases, 2 origin-binding proteins, 2 primases, 6 helicases, 6 DNA polymerases, 2 ligases and several other factors (Abu-Elneel et al., 2001; Downey et al., 2005; Lindsay et al., 2008; Liu et al., 2006; Saxowsky et al., 2003; Scocca and Shapiro, 2008)For example, TbPIF2 helicase and TbPRI1 primase have defined roles in maxicircle replication (Hines and Ray, 2010; Liu et al., 2009), while TbPIF1 and TbPRI2 have roles in minicircle replication (Hines and Ray, 2011; Liu et al., 2010). Three Family A DNA polymerase paralogs are required for *T. brucei* growth and kDNA maintenance in both life cycle stages (PCF and BSF): POLIB, POLIC, and POLID (Bruhn et al., 2010; Bruhn et al., 2011; Chandler et al., 2008; Klingbeil et al., 2002). While POLIB and POLID have demonstrated roles in minicircle replication, the precise role for POLIC in kDNA maintenance has not been determined.

Approximately 30 kDNA replication proteins have been characterized at the single protein level and localize to specific sites surrounding the kDNA disk including the KFZ, the AS flanking the disk, or within the disk. Given the unique spatiotemporal replication mechanism of the network, several studies have recently uncovered a complex choreography of kDNA replication proteins that likely coordinate subsets of proteins to specific sites during cell cycle progression (Abu-Elneel et al., 2001; Concepción-Acevedo et al., 2012; Concepción-Acevedo et al., 2018; Johnson and Englund, 1998; Peña-Diaz et al., 2017). POLID is the most dramatic example of cell cycle localization in which the protein colocalized with replicating minicircles at the AS only during kDNA S phase (1N1K*), and localized in the mitochondrial matrix during all other cell cycle stages (Concepción-Acevedo et al., 2012). Cell cycle stages are identified based on the single unit structures the nucleus (N) and kDNA (K), thus 1N1K are cells in G1, 1N1K* are cells undergoing kDNA S phase, 1N2K cells have segregated two progeny networks and are completing nuclear S phase and mitosis, and 2N2K cells have not yet undergone cytokinesis (2N2K). Another kDNA polymerase, POLIC, is initially detected in the KFZ/ULF, then accumulates at the AS only during kDNA S phase colocalizing with a fraction of POLID, thus supporting the idea of highly transient kDNA interactions (Concepción-Acevedo et al., 2018). The majority of the POLIC signal did not overlap with TAC102 signal. Additionally, when both proteins localized to the KFZ/ULF region, POLIC was more proximal to the kDNA than TAC102 (Concepción-Acevedo et al., 2018). What regulates the timing and recruitment associated with the protein choreography remains unknown.

The minicircle kDNA polymerases POLIB and POLID contain both a predicted C-terminal Family A DNA polymerase domain (POLA) and a 3’-5’ exonuclease domain. In contrast, POLIC is a 1649 aa protein (180 kDa) with only a conserved C-terminal POLA domain (315 aa). The remainder of this large protein is an uncharacterized N-terminal region we termed the UCR. Gene silencing of *POLIC* resulted in loss of fitness (LOF), a decrease in both minicircle and maxicircle copy number, and progressive loss of the kDNA network; hallmarks of a kDNA replication defect (Bruhn et al., 2010; Chandler et al., 2008; Liu et al., 2010). However, its silencing also resulted in cells displaying a predominant “small” kDNA phenotype rather than complete loss, an increase in the free minicircle population, and the appearance of ancillary kDNA, a phenotype associated with kDNA segregation defects (Archer et al., 2009; Clayton et al., 2011; Grewal et al., 2016; Trikin et al., 2016; Týč et al., 2015).

Therefore, the aim of this study was to apply a genetic complementation approach to dissect the pleiotropic RNAi phenotypes and further define the function of POLIC in kDNA maintenance. We present data that POLIC has two functional domains that contribute to kDNA maintenance; one for nucleotidyl incorporation and another that facilitates cell cycle localization and segregation of the kDNA network. An inactive POLA domain causes a dominant negative effect and exacerbates the kDNA replication defect of *POLIC* RNAi. Interestingly the UCR domain alone can restore localization to the AS during kDNA S phase. Both domains appear to contribute to proper segregation of kDNA. To our knowledge this is the first time a DNA polymerase is implicated in segregation of DNA and provides the strongest evidence thus far for interactions between the kDNA replication and segregation machineries.

## RESULTS

### *POLIC* 3′UTR RNAi and rescue with epitope-tagged wild-type protein

Previously, *POLIC* was silenced using an intermolecular dsRNA trigger targeting the open reading frame that led to the distinct pleiotropic small kDNA and ancillary kDNA phenotypes (Klingbeil, Motyka, Englund, 2002). To examine whether the POLA and UCR domains (Fig 1A) are necessary for either kDNA replication or segregation, we used a method that couples inducible RNAi-mediated gene silencing against the *POLIC* mRNA 3’ UTR with concurrent inducible ectopic expression of epitope-tagged variants in a structure-function analysis. We first tested whether *POLIC* 3’-UTR (IC-UTR) RNAi is sufficient to deplete *POLIC* mRNA. To generate the IC-UTR cell line, a 517 bp fragment immediately downstream of *POLIC* coding sequence was cloned into a stem-loop vector and subsequently introduced into the 29-13 cells (Wirtz, Leal, Ochatt, Cross, 1999). Tetracycline induction of dsRNA synthesis resulted in LOF following 10 doublings compared to uninduced cells (Fig 1A). RT-qPCR revealed a 65% reduction of *POLIC* mRNA following 48 hr of induction and the two paralogs (*POLIB* and *POLID*) as well as the downstream gene glutathione synthetase (Tb927.7.4000) exhibited no significant changes (Fig 1B). DAPI-staining of uninduced and induced cells was used to assess the effects of *POLIC 3’ UTR* silencing on kDNA networks. More than 300 parasites per time point were classified as possessing normal-size networks, small kDNA (networks unambiguously less than one-half the size of normal-sized networks seen in uninduced cells), or no kDNA if no extranuclear DAPI staining was observed despite viewing multiple focal planes. Using the previously reported categories, ten days of IC-UTR silencing resulted in cells that displayed small kDNA (74%), fragmented/ asymmetric networks (10.4%), normal kDNA (7.9%), and no kDNA (7.8%) (Fig 1C, E). These percentages closely resembled those from *POLIC* ORF silencing (Klingbeil, Motyka, Englund, 2002).

**Fig 1.**
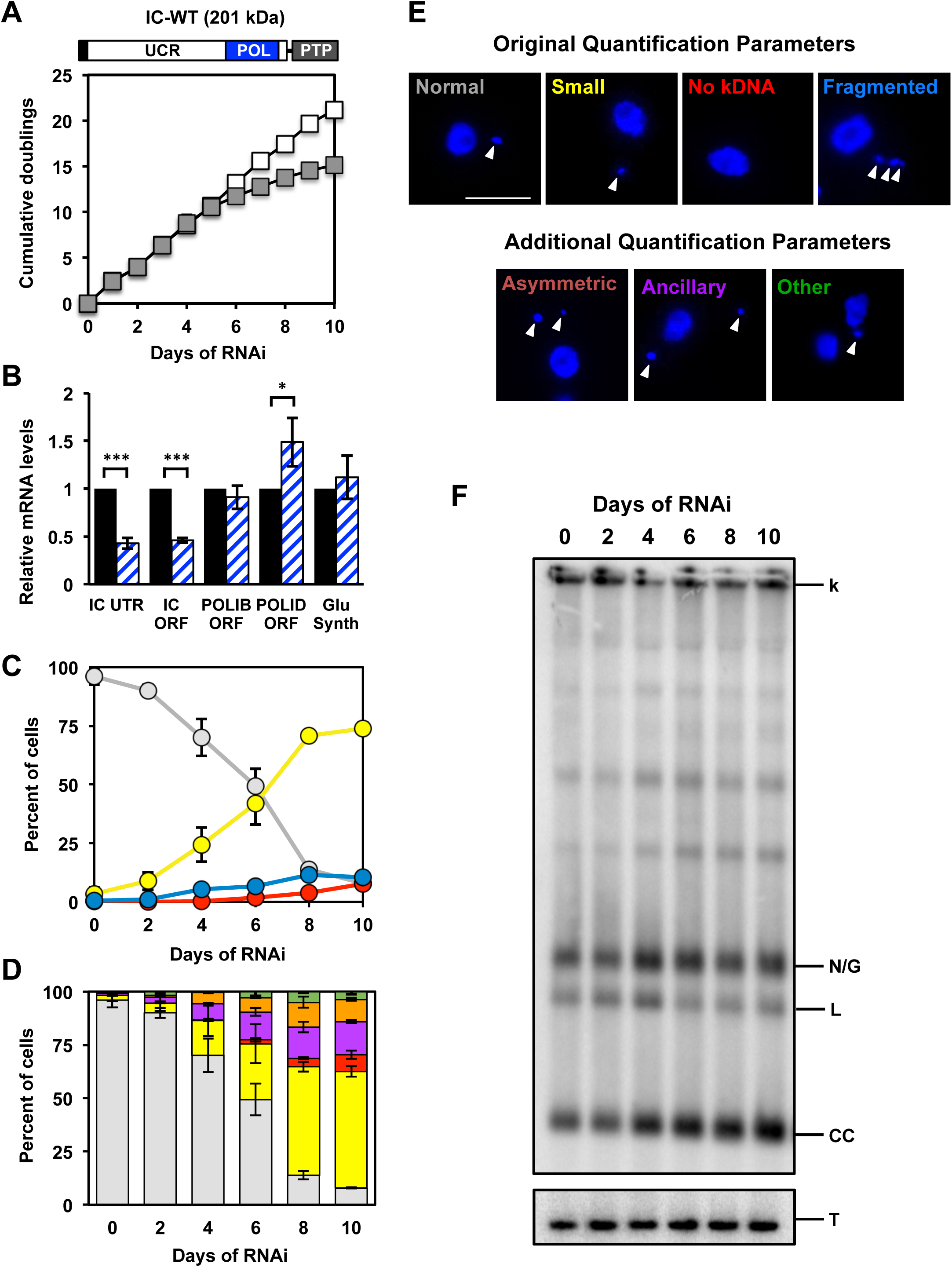
Phenotype of *POLIC* 3’UTR-targeted depletion. **(A)** Schematic of POLIC. Mitochondrial targeting sequence (black box), uncharacterized region (UCR), conserved Family A DNA polymerase domain (POL), and epitope tag (PTP). Growth curves of uninduced (open squares) and tetracycline-RNAi induced (gray squares) procyclic cells over 10 days. Error bars represent the standard deviation from the mean of three biological replicates. Above the growth curve, a schematic representation of full-length POLIC with the C-terminal PTP tag highlighting the UCR (white) and POL (blue) domains and tag (gray). **(B)** RT-qPCR analysis following 48 hours of uninduced (black) and tetracycline induced (blue/white hashed) IC-UTR cells. Data are normalized to *TERT* and error bars represent the standard deviation from the mean of three biological replicates. IC UTR, *POLIC* 3’ untranslated region; IC ORF, *POLIC* open reading frame; IB ORF, *POLIB* open reading frame; ID ORF, *POLID* open reading frame; Glu Synth, *Glutathione Synthetase*. Asterisks represent statistical significance based on pairwise Student’s T-test. * p < 0.1; ** p < 0.01; *** p < 0.001. **(C)** Quantification of kDNA morphology over the course of the induction by DAPI staining utilizing categories from Klingbeil, 2002. Over 300 cells were scored at each time point and error bars represent standard deviation from the mean of three biological replicates. gray, normal kDNA; yellow, small kDNA; red, no kDNA; blue, fragmented kDNA. **(D)** Quantification of kDNA morphology over the course of the induction by DAPI staining utilizing new parameters. Over 300 cells were scored at each time point and error bars represent standard deviation from the mean of three biological replicates. Gray, normal kDNA; yellow, small kDNA; red, no kDNA; purple, ancillary kDNA; orange, asymmetric size; green, other. **(E)** Representative images of DAPI stained cells for the various kDNA phenotypes. Scale bar, 5 µm. **(F)** Representative Southern blot showing the effect of IC 3’ UTR silencing on the free minicircle population. k, kDNA network; N/G, nicked/gapped; L, linearized; CC, covalently closed; T, tubulin.

To further dissect the multiple kDNA-associated defects, cells that contained ancillary or asymmetrically divided kDNA were also quantified. Ancillary kDNA and asymmetric network division are hallmarks of segregation defects (Grewal et al., 2016; Trikin et al., 2016; Zhao et al., 2008).By further subdividing the categories, IC-UTR RNAi still produced cells with a majority of small kDNA (54.7%) vs normal kDNA (7.9%), no kDNA (7.5%) and other (3.7%), but also highlighted the segregation defect signatures of ancillary kDNA (15.6%), asymmetric division (10.4%), (Fig 1D, E). Based on the proximity of POLIC and TAC102 in the KFZ/ULF, we wanted to test whether TAC102 localization is impacted during *POLIC* depletion. After 4 days of *POLIC* depletion, the majority of TAC102 localized properly in close proximity to the basal body while some cells showed TAC102 co-localizing with ancillary kDNA as previously reported (Trikin et al., 2016). TAC102 protein appeared stable throughout the induction. (Fig S1). Lastly, by analyzing the free minicircle population using Southern blotting and a minicircle-specific probe, both unreplicated CC monomers and newly replicated N/G molecules increased when *POLIC* was depleted compared to the uninduced control, another indication of a defect not strictly related to minicircle replication (Fig 1F). Taken together, IC-UTR silencing phenocopied the previously reported LOF, and kDNA replication defect but also displayed clear signatures of a segregation defect.

To determine whether the deficiencies caused by IC-UTR RNAi could be rescued by ectopic expression of PTP-tagged wild-type POLIC (IC-WT, 201 kDa), we transfected the IC-UTR cells with pLewIC-WT-PTP^Puro^ (RNAi+OE). IC-WT complementation alleviated the RNAi-mediated defects with consistent expression of IC-WT throughout the induction (Fig S2A-E). IC-WT levels were four-fold above endogenous POLIC protein detected in a cell line engineered to endogenously express the epitope-tagged POLIC from a single allele (+C, Fig S2C) (Concepción-Acevedo, Miller, Boucher, Klingbeil, 2018). Additionally, we analyzed the cell cycle localization of IC-WT and found discrete antipodal POLIC-PTP foci formation in 35% of an unsynchronized population almost exclusively in the 1N1K* stage, similar to endogenous 26% POLIC-PTP AS foci detected in the single expresser cell line (Fig S2F) (Concepción-Acevedo, Miller, Boucher, Klingbeil, 2018).

We also transfected pLewIC-WT-PTP^Puro^ into parental 29-13 cells generating the IC-WT overexpression (OE) cell line to study the effects of increased IC protein in the absence of *POLIC 3’UTR* RNAi. IC-WT overexpression (8-fold) for 10 days did not interfere with fitness or kDNA maintenance and IC-WT localization was similar to endogenous POLIC but IC-WT OE cells displayed additional matrix localization (Fig S3). The localization patterns from IC-WT overexpression and complementation were nearly identical (Fig S2F and S3C). Therefore, IC-UTR RNAi was specific based on the complete rescue using IC-WT and the unique pleiotropic phenotype could be attributed solely to silencing of *POLIC*.

### POLIC DNA polymerase activity is essential for kDNA replication and cell cycle localization

Mutational and structural analyses of Family A DNA polymerases have revealed two invariant Asp residues essential for catalytic activity that coordinate two divalent metal ions for transition state stabilization necessary for all nucleotidyl transferases (Steitz, 1998, Sousa 1996). POLIC and the other *T. brucei* mitochondrial DNA polymerases contain Family A conserved motifs A, B and C and the non-variant Asp residues (Klingbeil 2002). It is possible that the polymerase function may not be the essential function of POLIC. In yeast, the DNA polymerase and 3’-5’ exonuclease activity of DNA Pol ε are nonessential for cell growth, whereas the protein’s non-catalytic C-terminal domain is required (Kesti et al., 1999; Dua et al., 1999).

To address whether the nucleotidyl transferase activity is essential for POLIC *in vivo* functions, we generated a full-length POLIC PTP-tagged variant with catalytic residue mutations (D1380A, D1592A) to inactivate polymerase activity (IC-DEAD), and transfected it into the overexpression (OE) and the IC-UTR RNAi backgrounds (RNAi+OE). To confirm the mutated Asp residues successfully inactivated POLIC polymerase activity, IC-WT and IC-DEAD were overexpressed for 48 hours, immunoprecipitated and analyzed by western blot and an *in vitro* primer extension assay. The POLIC variants were detected at their predicted sizes (IC-DEAD, 201 kDa; IC-POLA, 75 kDa, IC-UCR, 160 kDa), but only IC-WT demonstrated nucleotidyl transferase activity (Fig S4).

Interestingly, IC-DEAD overexpression led to a dominant-negative LOF following just 4 days of induction. The number of cumulative doublings after 10 days of IC-DEAD overexpression was 13.7 (Fig 2A, left graph) compared to the 15.1 cumulative doublings following IC-UTR RNAi (Fig 1A). However, complementation with the IC-DEAD variant exacerbated the LOF and reduced the cumulative doublings to 9.1 after 10 days (Fig 2A, right graph). RT-qPCR confirmed that IC-DEAD complementation depleted endogenous *POLIC* transcripts, while abundance of the variant was significantly higher than endogenous *POLIC* (Fig 2B). The abundance of the paralog transcripts was also elevated compared to uninduced. IC-DEAD protein levels were initially high compared to endogenous POLIC in both backgrounds (OE, 16.6-fold; RNAi+OE, 12.5-fold) but were reduced to 3-fold by the end of the 10 day induction based on ImageJ analyses from Western blot data (Fig 2C). IC-DEAD-PTP signal was concentrated on or near the kDNA at all cell cycle stages in OE (98%) and RNAi+OE (99%) cell lines (Fig 2D-E) in striking contrast to endogenous POLIC-PTP and IC-WT-PTP foci found exclusively in the 1N1K* stage (compare Figs 2E and S2F). IC-DEAD complementation also occasionally displayed ancillary kDNA co-localized with TAC102 (Fig S1).

**Fig 2.**
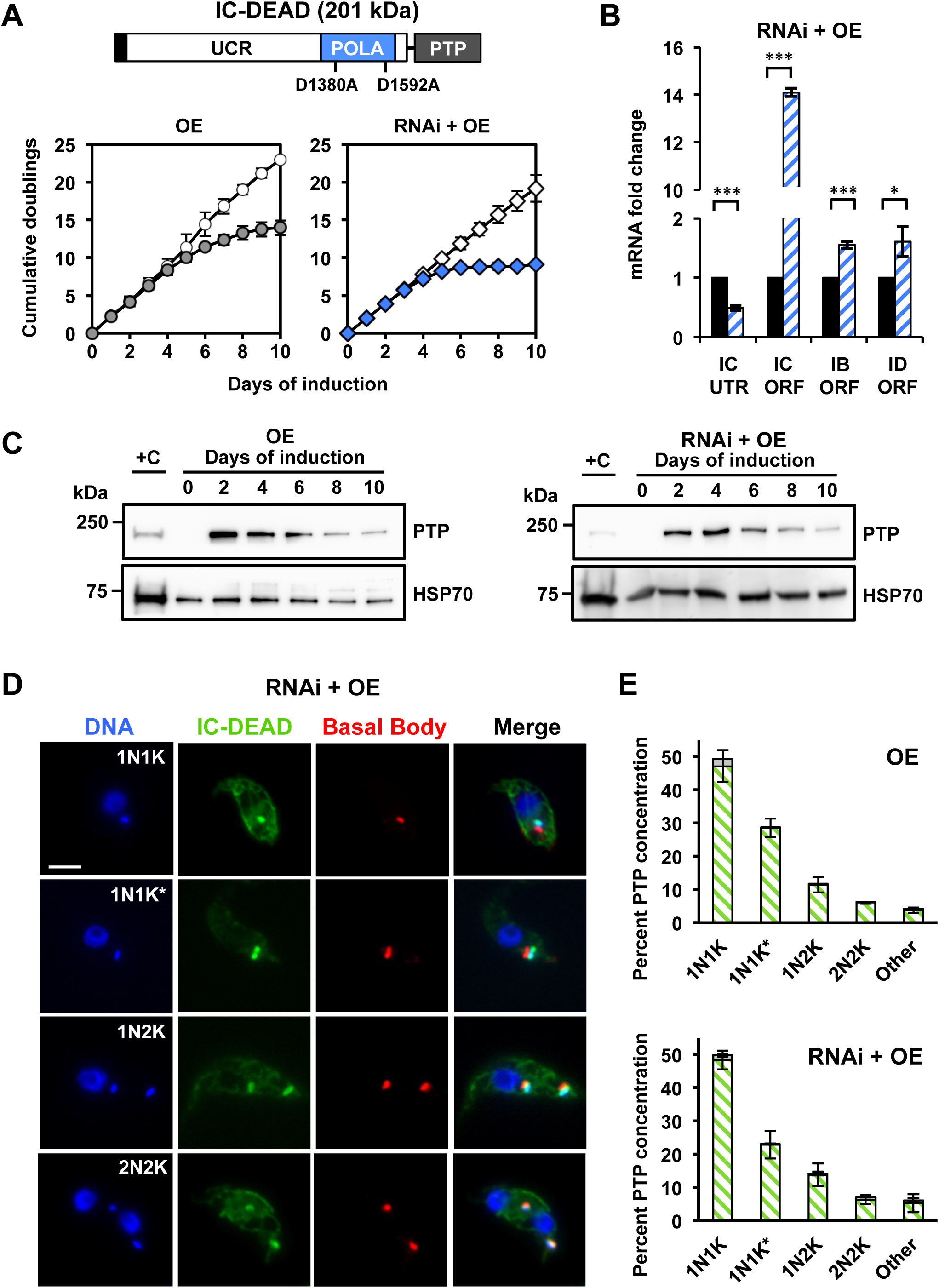
Overexpression of IC-DEAD results in a dominant-negative phenotype while complementation exacerbates loss of fitness. **(A)** POLIC domain structure. As describe in Figure 1A with the asp acid residues mutated to alanine, and growth curves of uninduced (OE, open circles; complementation, open diamonds) and tetracycline-induced (OE, gray circles; complementation, blue diamonds) procyclic cells over 10 days. Error bars represent the standard deviation from the mean of three biological replicates. **(B)** RT-qPCR analysis of uninduced (black) and induced (blue/white hashed) cells following 48 hours of tetracycline induction. Data are normalized to *TERT* and error bars represent the standard deviation from the mean of three biological replicates. IC UTR, *POLIC* 3’ untranslated region; IC ORF, *POLIC* open reading frame; IB ORF, *POLIB* open reading frame; ID ORF, *POLID* open reading frame. Asterisks represent statistical significance based on pairwise Student’s T-test. * p < 0.1; ** p < 0.01; *** p < 0.001. **(C)** Western blot detection of the PTP tag and HSP70 protein levels following 10 days of induction. 1×10^6^ cell equivalents was loaded into each well except for C where 5×10^6^ cell equivalents was loaded. +C, single expresser control cell line. **(D)** Representative images of IC-DEAD at each cell cycle stage. DAPI-staining (blue); anti-protein A (green); YL1/2 (red). Scale bar, 5 μm. **(E)** Quantification of PTP concentrations at each cell cycle stage in the overexpression (OE, top) and complementation (RNAi + OE, bottom) cell lines after 48 hours of induction. Percentages refer to the fraction of total cells in the population. Foci positive (Green/white hashed); foci negative (gray). Error bars represent the standard deviation from the mean of three biological replicates.

Next, we examined the effects of IC-DEAD variant on kDNA maintenance in both backgrounds (OE and RNAi+OE). Ten days of IC-DEAD overexpression in the absence of *POLIC* RNAi resulted in a predominantly small kDNA phenotype (43.5%), followed by no kDNA (26.5%), ancillary (14.3%), other (11%), and normal (4.7%) kDNA (Fig 3A). In striking contrast to both OE and the parental RNAi phenotypes, IC-DEAD complementation displayed a majority of cells with no kDNA (59.1%), as well as small (16.1%), ancillary (10.8%), other (9.6%), and normal (4.4%) kDNA that correlated with the exacerbated LOF (Fig 3D). The IC-DEAD variant causes a more rapid onset of aberrant kDNA (after 2 days) in both genetic backgrounds and shifts from a predominantly small kDNA to no kDNA for the IC-DEAD complementation (Figs 1D, 3A, 3D).

**Fig 3.**
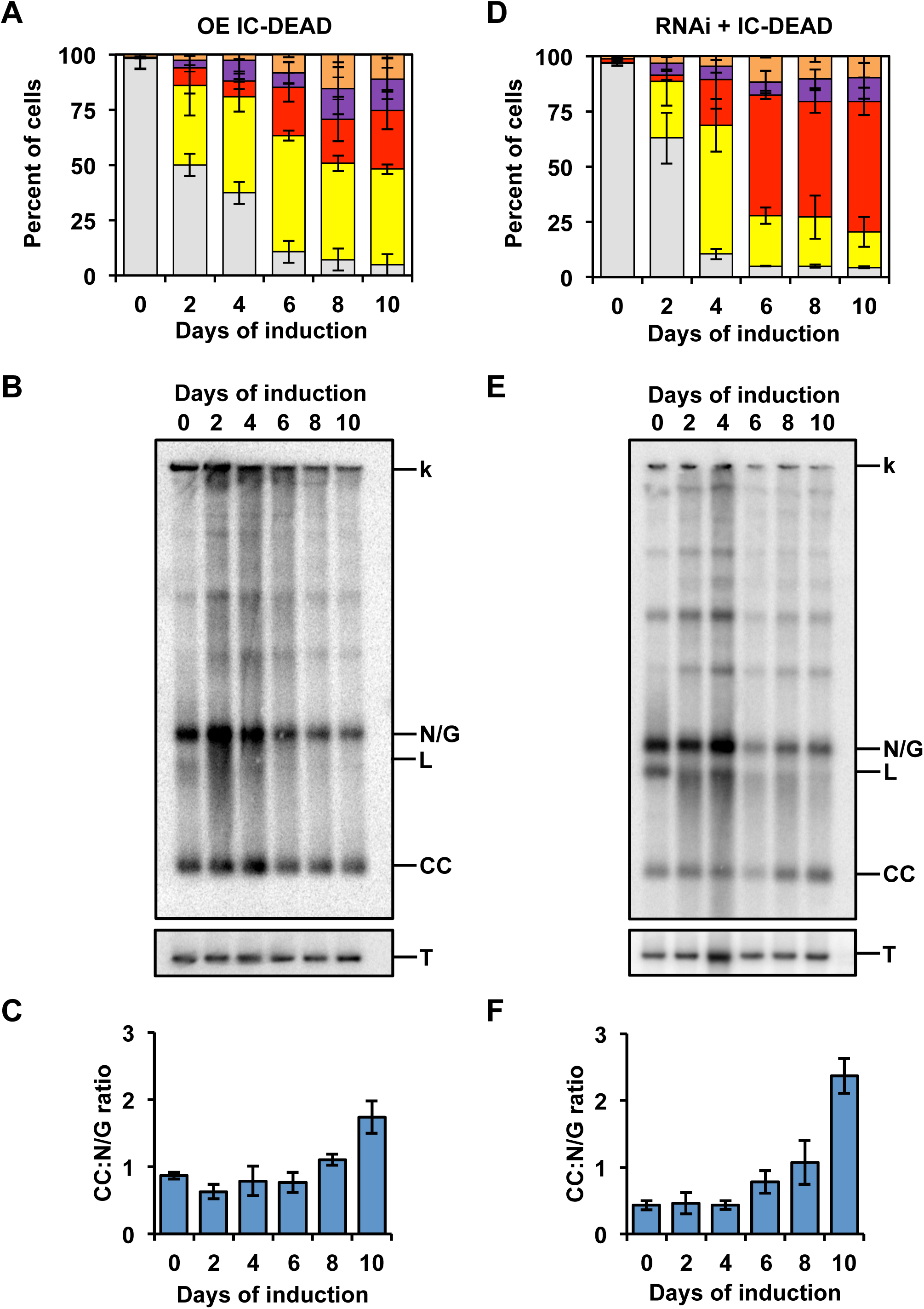
kDNA replication phenotype analysis of IC-DEAD overexpression and complementation. **(A)** Quantification of kDNA morphology for OE IC-DEAD over the course of the induction. Over 300 cells were scored at each time point and error bars represent standard deviation from the mean of three biological replicates. Gray, normal kDNA; yellow, small kDNA; red, no kDNA; purple, ancillary kDNA; orange, asymmetric size; green, other. **(B)** Representative Southern blot showing the effect on free minicircle population. k, kDNA network; N/G, nicked/gapped; L, linearized; CC, covalently closed; T, tubulin. **(C)** Ratio of CC:N/G over the course of the induction. Phosphoimager quantification was used to plot the relative abundance of unreplicated CC and newly replicated N/G intermediates. Error bars represent the standard deviation from the mean of three biological replicates. **(D-F)** Same as A-C but for the RNAi + IC-DEAD cell line.

In addition to progressive loss of the kDNA network, hallmarks of perturbing kDNA replication include a decrease in minicircle and maxicircle copy number as well as an increased ratio of unreplicated covalently closed (CC) to nicked/gapped (N/G) progeny (Bruhn et al., 2010; Liu et al., 2010; Scocca and Shapiro, 2008). Both IC-DEAD overexpression and complementation led to a progressive decrease in free minicircle species (Fig 3B, E). In both genetic backgrounds, the free minicircles initially increased (Days 2, 4) when the IC-DEAD variant was present. During the later days of the inductions, the abundance of CC and N/G minicircles quickly declined, which resembled the defect caused by RNAi of the paralog POLID (Chandler et al., 2008). When comparing unreplicated CC to replicated N/G species, IC-DEAD overexpression showed a 1.7-fold increase in the CC:N/G ratio and complementation displayed a 5.5-fold increase at day 10 compared to uninduced cells (Fig 3C, F).

These data indicate that IC-DEAD could not rescue any of the pleiotropic effects from *POLIC* RNAi. Additionally, the variant resulted in disruption of cell cycle localization and displayed an overt kDNA replication defect with a decrease in N/G progeny suggesting that the nucleotidyl incorporation is essential for POLIC function.

### The POLA domain rescues minicircle replication defects

Next, we determined if the conserved POLA domain alone was sufficient to restore POLIC function *in vivo*. We generated an N-terminal truncated POLIC PTP-tagged variant (IC-POLA, 437 aa), and transfected it into the overexpression (OE) and the IC-UTR RNAi backgrounds (RNAi+OE). Consistent IC-POLA overexpression (5.4-fold compared to endogenous) did not affect parasite fitness (Figs 4A, Fig S5A) or kDNA maintenance determined by both DAPI staining which showed >90% normal cells at day 10, and by Southern blotting which indicated no perturbation of free minicircle levels (Fig S5B, C). Surprisingly, immunoprecipitated IC-POLA did not display nucleotidyl incorporation activity in the primer extension assay (Fig S4).

**Fig 4.**
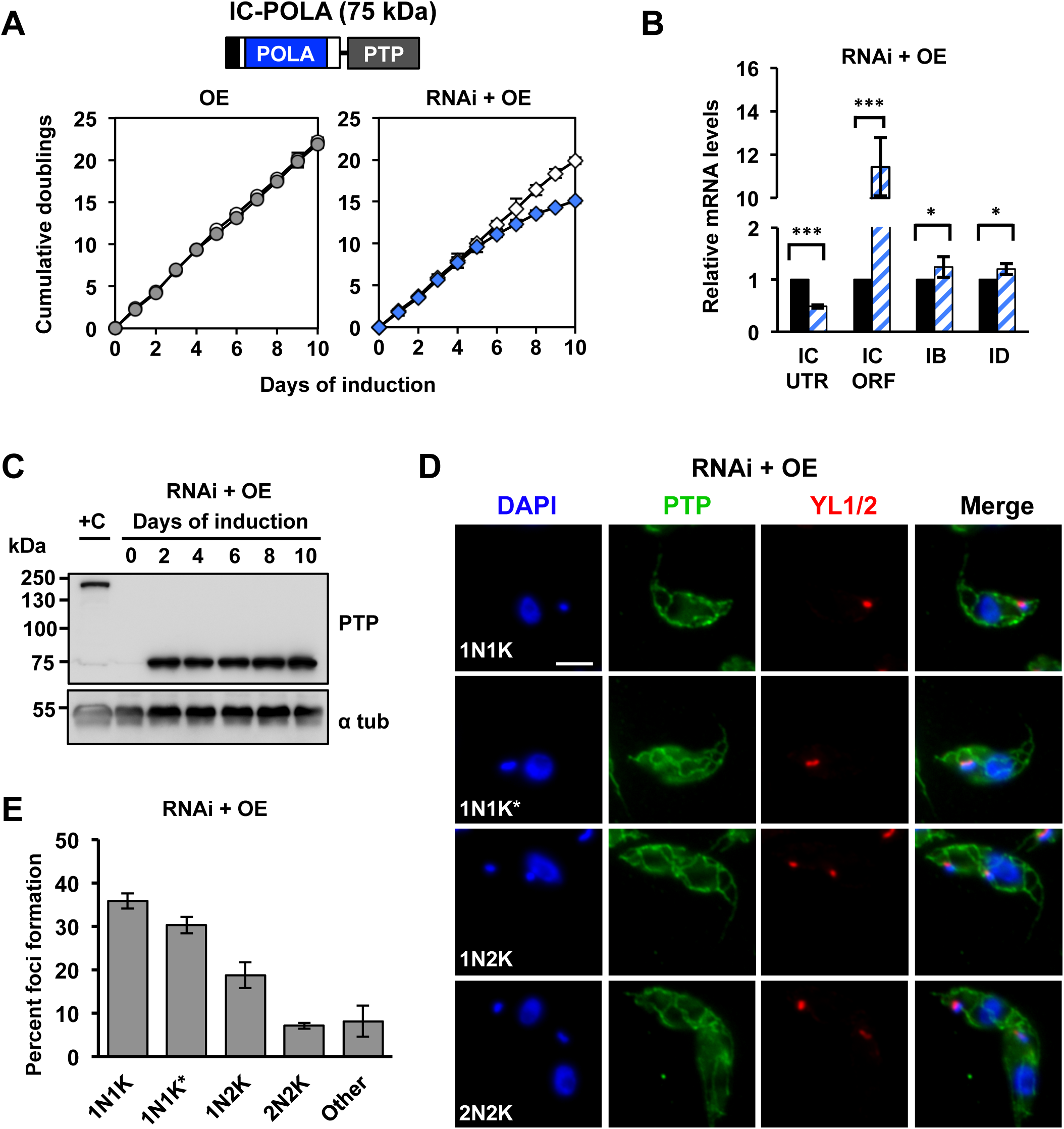
IC-POLA complementation is not sufficient to rescue the RNAi phenotype. (A) Growth curves of uninduced (OE, open circles; complementation, open diamonds) and tetracycline-induced (OE, gray circles; complementation, blue diamonds) cell cultures over 10 days of IC-POLA in the overexpression or complementation cell lines. (B) RT-qPCR analysis of uninduced (black) and induced (blue/white hashed) cells following 48 hours of tetracycline induction. Data are normalized to *TERT* and error bars represent the standard deviation from the mean of three biological replicates. IC UTR, *POLIC* 3’ untranslated region; IC ORF, *POLIC* open reading frame; IB, *POLIB*; ID, *POLID*. Asterisks represent statistical significance based on pairwise Student’s T-test. * p < 0.1; ** p < 0.01; *** p < 0.001. (C) Western blot detection of the PTP tag and alpha tubulin protein levels following 10 days of induction. (D) Representative images of IC-POLA at each cell cycle stage. Scale bar, 5 μm. (E) Quantification of foci formation at each cell cycle stage. Green, foci positive; gray, foci negative.

IC-POLA complementation closely resembled the parental IC-UTR LOF (15.1 cumulative doublings, Day 10) with robust knockdown of endogenous *POLIC* mRNA (Fig 4A, B). Western blot data showed consistent protein expression for 10 days (4.1-fold higher than endogenous) (Fig 4C). IC-POLA showed only mitochondrial matrix localization and weak signal near the kDNA regardless of cell cycle stage, and no antipodal PTP foci were ever detected (Fig 4D, E). IC-POLA complementation displayed cells with small (46.7%), no (9.8%), normal (7.5%), and other (3.1%) kDNA (Fig 5A). Interestingly, IC-POLA cells with ancillary (19.5%) or asymmetric (13.3%) kDNA were elevated compared to the combined percentage of fragmented/asymmetric (25.5%) kDNA for the parental IC-UTR RNAi defect, suggesting that the polymerase domain alone could not rescue the segregation defect. TAC102 was detected co-localizing with a portion of the ancillary kDNA (Fig S1). Furthermore, in the presence of the IC-POLA variant, there was no detectable accumulation of free minicircles over the course of the induction (Fig 5B, C), and there were no statistically significant changes in the CC:N/G ratio across the 10 day induction demonstrating that POLA domain could rescue the minicircle replication defect.

**Fig 5.**
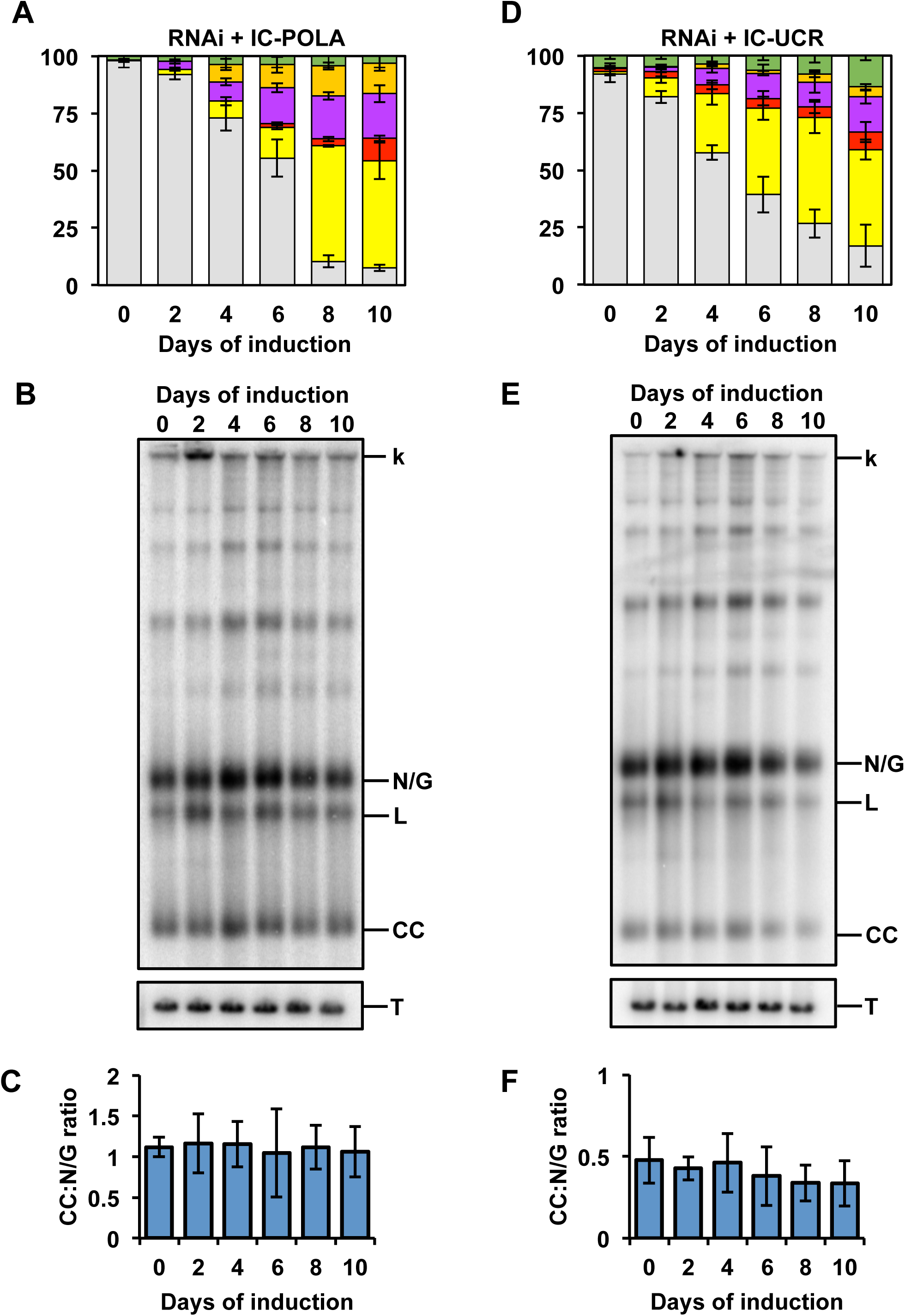
kDNA replication phenotype analysis of IC-POLA and IC-UCR complementation. **(A)** Quantification of RNAi + IC-POLA kDNA morphology over the course of the induction. Over 300 cells were scored at each time point and error bars represent standard deviation from the mean of three biological replicates. Gray, normal kDNA; yellow, small kDNA; red, no kDNA; purple, ancillary kDNA; orange, asymmetric size; green, other. **(B)** Representative Southern blot showing the effect on free minicircles of RNAi + IC-POLA. k, kDNA network; L, linearized; N/G, nicked/gapped; CC, covalently closed; T, tubulin. **(C)** Quantification of the CC:N/G ratio over the course of the induction. Phosphoimager quantification was used to plot the relative abundance of unreplicated CC and newly replicated N/G intermediates. Error bars represent the standard deviation from the mean of three biological replicates. **(D-F)** Same as A-C but for the RNAi + IC-UCR cell line.

Collectively, these data show that the C-terminal polymerase domain alone could restore the free minicircle population even though it could not properly localize to the AS. Additionally, the increased segregation defect suggested that the UCR is necessary for kDNA segregation and cell cycle dependent localization.

### The UCR domain partially rescues IC-UTR RNAi defects

The N-terminal UCR (1291 amino acids) of POLIC has no conserved domains or motifs but does contain two hydrophobic patches and arginine methylation sites (R420, R1250, R1260) that were detected in the *T. brucei* mitochondrial arginine methylproteome (Fisk et al 2012). To evaluate a role in kDNA segregation, we generated a POLIC variant that contained only the UCR region (IC-UCR), and transfected it into the overexpression (OE) and the IC-UTR RNAi backgrounds (RNAi+OE). IC-UCR overexpression was consistent across the induction (5.3-fold increase compared to endogenous), did not affect parasite fitness, and no kDNA replication defects were detected (Fig 6A, S4D-F).

**Fig 6.**
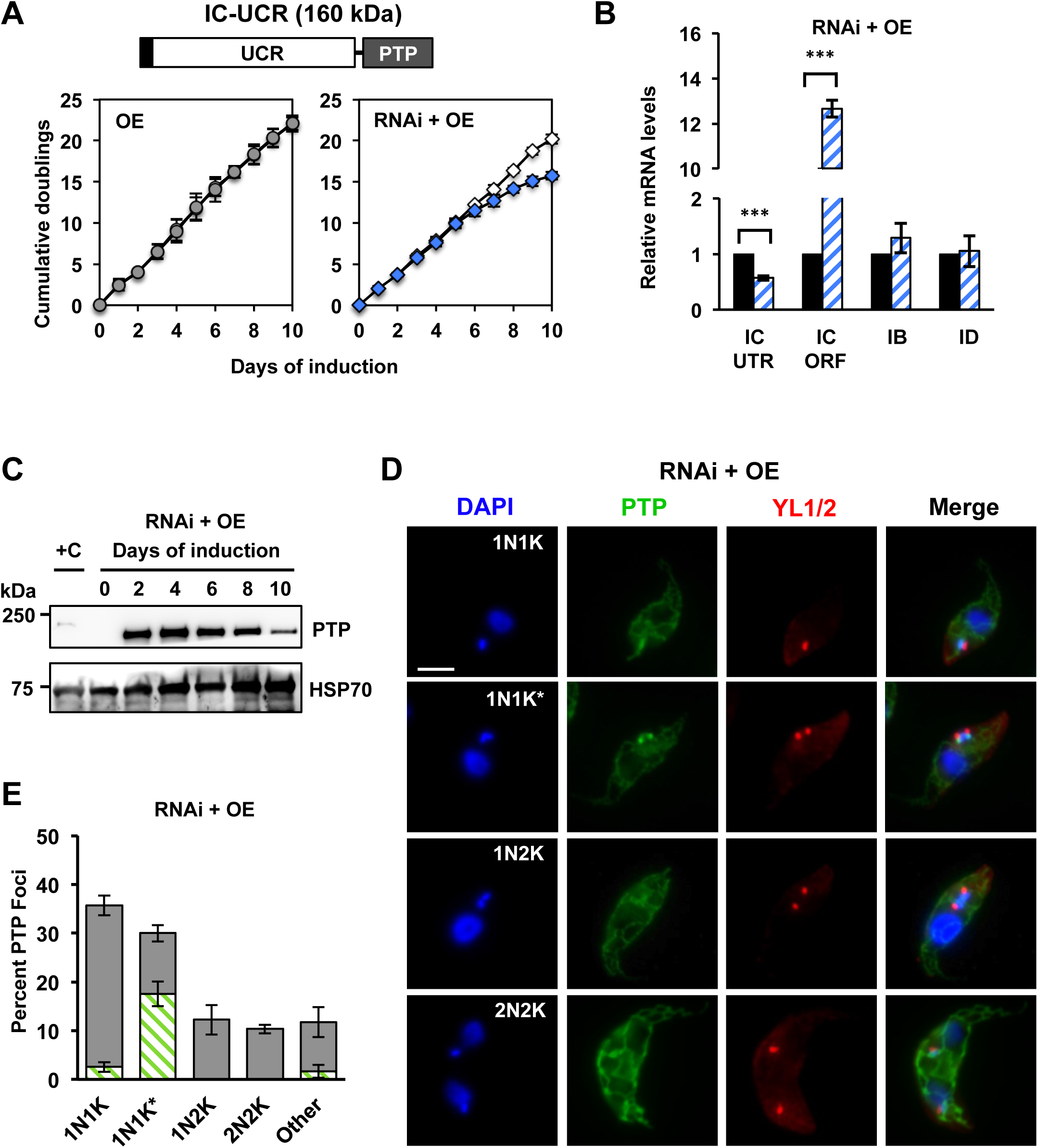
IC-UCR complementation is not sufficient to rescue the RNAi phenotype. **(A)** Growth curves of uninduced (OE, open circles; complementation open diamonds) and tetracycline-induced (OE, gray circles; complementation blue diamonds) cell cultures over 10 days of IC-UCR in the overexpression or complementation cell lines. Above the growth curve, a schematic representation of IC-UCR variant with the C-terminal PTP tag highlighting the UCR (white) domain and tag (gray). **(B)** RT-qPCR analysis of uninduced (black) and induced (blue/white hashed) cells following 48 hours of tetracycline induction. Data are normalized to *TERT* and error bars represent the standard deviation from the mean of three biological replicates. IC UTR, *POLIC* 3’ untranslated region; IC ORF, *POLIC* open reading frame; IB, *POLIB*; ID, *POLID*. **(C)** Western blot detection of the PTP tag and HSP70 protein levels following 10 days of induction. **(D)** Representative images of IC-UCR at each cell cycle stage. Scale bar, 5 μm. **(E)** Quantification of foci formation at each cell cycle stage. Percentages refer to the fraction of total cells in the population. Green/white hashed, foci positive; gray, foci negative. Error bars represent the standard deviation from the mean of three biological replicates.

IC-UCR complementation resulted in LOF with 15.7 cumulative doublings after 10 days of induction (Fig 6A). RT-qPCR displayed robust *POLIC* knockdown and Western blot data revealed initially high protein levels (18-fold increase over endogenous) that continually decreased over 10 days (4-fold) (Fig 6B, C). There was clear formation of IC-UCR-PTP foci at AS in 17.6% of 1N1K* cells compared to IC-WT complementation (30%) and endogenous localization (26%) and IC-UCR also displayed weak signal near the kDNA at other cell cycle stages (Fig 6D, E).

IC-UCR complementation displayed mostly cells with small (42%), normal (17%), ancillary (15.3%), other (13.7%), no (7.7%), and asymmetric (4.3%) kDNA (Fig 5D). Interestingly, the IC-UCR variant resulted in more cells with normal kDNA (17 vs. 7.8%), and fewer cells with segregation defects when compared to IC-UTR RNAi alone (19.6 vs 25.5%) indicating a partial rescue of the segregation phenotype. Despite the reduction in cells displaying asymmetric networks, the percentage of cells displaying ancillary kDNA did not significantly improve and TAC102 still co-localized with ancillary kDNA (Fig S1). In the presence of the IC-UCR variant, there were no statistically significant changes in the CC:N/G ratio even though there was a decrease in free minicircles after 8 days of complementation (Fig 5E, F). Collectively, these data revealed POLIC has distinct functions in both kDNA replication and segregation and that the UCR mediates antipodal site localization and the segregation function.

## DISCUSSION

Trypanosome kDNA replication presents an interesting model system in which replication of a single nucleoid (the kDNA network) occurs once every cell cycle and segregation of daughter networks is mediated by a structure linked to the flagellar basal body called the TAC. The current kDNA replication model indicates spatial and temporal separation for early events occurring in the KFZ and later Okazaki processing at the AS. Additionally, three Family A DNA polymerases (Pol) are essential for kDNA replication. POLIB and POLID have demonstrated roles in minicircle replication, but the precise role for POLIC in kDNA maintenance was unclear. Although the precise functions performed by the paralogs on minicircles and maxicircles have not yet been defined, the trio of Pols superficially resembles the replicative polymerases Pol α, δ, and ε required for nuclear DNA replication. Understanding catalytic and non-catalytic roles may provide insight as to why trypanosomes utilize three distinct Pols for kDNA replication.

POLIC shares no similarity with other proteins outside of the conserved POLA domain at the C-terminus. Consistent with an essential function in kDNA maintenance, POLIC and the overall protein structure is conserved in all of the 41 currently sequenced trypanosomatid organisms and some free-living relatives available at TriTrypDB.org (Aslett et al., 2010). To further understand the role of POLIC in kDNA replication, we investigated the structure–function relationship by ectopically expressing several POLIC variants in trypanosome cells that are deficient in endogenous POLIC. Tetracycline-inducible expression of a dsRNA targeting the 3’-UTR of a gene and co-expression of an ectopic copy of the RNAi target is a validated approach for *in vivo* complementation studies on flagellar motility, cytokinesis, mitochondrial protein import, TAC function, and transferrin receptor trafficking in *T. brucei* (Käser et al., 2016; Käser et al., 2017; Ralston et al., 2011; Trikin et al., 2016; Weems et al., 2015; Yu et al., 2012). By using this approach we demonstrated that WT-IC could rescue the replication and segregation defects and we defined two functional domains: one for nucleotidyl incorporation and a non-catalytic domain for AS localization and kDNA segregation. This dual-functioning protein provides direct evidence for interaction between the replication and segregation machineries.

How the POLIC polymerase domain participates at replication forks remains an open question. Although the immunoprecipitated POLA domain did not have detectable activity in a primer-extension assay (Fig. S4), IC-POLA complementation indicated that the polymerase domain alone is sufficient to support minicircle replication even though IC-POLA localized mainly to the mitochondrial matrix (Fig. 4,5B, C). IC-POLA includes all the conserved Family A motifs, but this truncated fragment may lack important upstream residues required for binding the primer-template in the *in vitro* assay. Despite IC-POLA mislocalization during complementation, abundance was likely high enough to support the polymerase function. It is compelling that the IC-DEAD variant defective in nucletotidyl incorporation generated the most striking kDNA replication defects in both overexpression and complementation. IC-DEAD overexpression caused a dominant negative LOF with a kDNA replication defect (Fig. 2A), and abolished POLIC cell cycle localization. Instead of mitochondrial matrix localization, IC-DEAD always localized with or around the kDNA disk (Fig. 2D, E). Interestingly, there are now 250 pathogenic mutations in human mitochondrial DNA polymerase gamma that interfere with dNTP selectivity, processivity or accessory subunit binding with many of the dominant mutations located in the polymerase domain that result in loss of mtDNA (Stumpf et al., 2013). It is possible that IC-DEAD was bound to minicircles and/or maxicircles outcompeting endogenous POLIC for substrate, inhibited access of other replication proteins, or sequestered replication factors. IC-DEAD complementation results support the idea that access of other replication proteins was impacted. The dramatic loss of kDNA (Fig. 3D) and increase in unreplicated free minicircles (Fig. 3E,F) are hallmarks of a kDNA replication defect that closely resemble depletion of POLID, a known minicircle replication factor that localizes to AS only during kDNA S phase (Chandler et al., 2008; Concepción-Acevedo et al., 2012). In addition to masking the original small kDNA phenotype, only the IC-DEAD variant increased transcript abundance of POLIB and POLID suggesting functional interplay among the paralogs (Fig. 2B). We previously showed that POLID depletion disrupted localization of POLIC to AS (Concepcion et al 2018). However, the mechanisms(s) governing AS localization and paralog interactions were never determined.

Despite lacking any recognizable catalytic domain, the UCR appears to contain a determinant for localization to AS. IC-UCR complementation resulted in PTP foci formation that closely resembled the localization pattern for endogenous POLIC where signal was detected in the KFZ in 1N1K cells, IC-UCR-PTP foci at AS were detected in 1N1K* cells, and the signal was disperse in the other cell cycle stages (Fig. 6D, E). Although two different methods were used to evaluate the percentage of cells in kDNA S phase, the proportion of PTP foci positive cells in this stage were equivalent (0.59) (Concepcion et al 2018). Cell cycle localization of POLIC may be a mechanism that discriminates between the two functional domains of this protein. For example, nucleotidyl incorporation activity may be utilized when localized in the KFZ, while the segregation function may be more important at AS. The IC-UCR variant contains three residues that are arginine methylated (Fisk et al 2012). Arginine methylation is a reversible post-translational modification with regulatory roles that may contribute to POLIC localization at AS through proteins-protein interactions. IC-UCR is the first sequence from a kDNA protein shown to promote AS localization and further experimentation is necessary to refine the specific sequences determinants involved.

Another non-catalytic role for the UCR is associated with kDNA segregation. IC-UCR complementation partially rescued RNAi by reducing the number of cells with asymmetrically sized networks and increasing the number of cells with normal kDNA (Fig. 5D). The decline in IC-UCR abundance over the course of the complementation provides one explanation for the incomplete rescue. Alternatively, the POLA domain may also have a role in segregation through protein interactions. Surprisingly, no POLIC variant complementation was able to reduce the appearance of ancillary kDNA (Fig. 3D, 5A, D) suggesting that a fully functional protein is critical to suppress the ancillary kDNA phenotype. It is likely that POLIC is not playing a direct role in segregation but is impacting other proteins essential for segregation.

POLIC is not a classical TAC component based on the criteria outlined by Schneider and Ochsenreiter (2018). IC-UTR RNAi does not result in 1 cell with a large kDNA and the other lacking kDNA. Instead, there is unequal partitioning of a reduced sized network due to the POLIC replication defect. Additionally, ancillary kDNA is an unusual aberrant phenotype that appears when elements related to the mitochondrial membrane or the TAC are perturbed as seen with functional disruptions of PUF9, ACP, PNT1, HSP70/HSP40, and TAC102 (Archer et al., 2009; Clayton et al., 2011; Grewal et al., 2016; Trikin et al., 2016; Týč et al., 2015). The TAC component most proximal to the kDNA is TAC102. When the C-terminus of this protein is obstructed, TAC102 cannot connect to the upstream TAC components properly (and subsequently to the basal body) resulting in mislocalization with ancillary kDNA (Trikin et al 2016). It is interesting that complementation with all POLIC variants produced ancillary kDNA with mislocalization of TAC102 independent of their rescue status (Fig S1). POLIC is more proximal to the kDNA than TAC102 (Concepcion et al 2018) and our data suggest that a fully functional POLIC is important for TAC102 to connect to upstream TAC components. At this time we do not know if the TAC is required for proper localization of POLIC or how POLIC might interact with the TAC. The recently described minicircle reattachment protein MiRF172 may provide a bridge between POLIC and TAC102 since MiRF172 remains associated with isolated flagella following biochemical fractionations, and its localization is more proximal to the kDNA than TAC102 (Amodeo, Jakob, Ochsenreiter, 2018). The proximity of the replication and segregation machinery in *T. brucei* could suggest a physical interaction of the components involved in the two processes. Thus far, the dual-functioning DNA polymerase, POLIC, is the clearest example of a protein bridging these two essential processes.

## MATERIALS AND METHODS

### For Primer sequences refer to Supplemental Table 1

#### DNA constructs

##### (i) RNAi

A pStL (stem-loop) vector for inducible gene silencing of *POLIC* was constructed as previously described (Wang, Morris, Drew, Englund, 2000). Briefly, 517 bp of *POLIC* 3’ UTR sequence was PCR amplified from TREU 927 gDNA using primers listed in Supplemental Table 1 to generate the two fragments for subsequent cloning steps, The resulting vector, pStLIC3’UTR was EcoRV-linearized for genome integration.

##### (ii) Site-Directed Mutagenesis

Full-length wild-type *POLIC* (Tb927.7.3990) was PCR amplified from TREU 927 gDNA using Phusion DNA polymerase and subcloned into the pCR®-Blunt II-TOPO vector (Invitrogen). Mutation of 1380Asp and 1592Asp to Ala was performed by site-directed mutagenesis using the QuikChange Lightning Multi Site-Directed Mutagenesis Kit (Agilent) generating pTOPO-IC-DEAD.

##### (iii) Inducible expression of POLIC variants

To create pLew100-PTP^Puro^ for inducible expression of PTP tagged POLIC variants, the Myc tag of pLew100-Myc^Puro^ (gift from Dr. Laurie Read, SUNY Buffalo) was replaced with the PTP tag. Briefly, the PTP tag was PCR amplified from pC-PTP-NEO (Schimanski et al 2005) and cloned into pCR®-Blunt II-TOPO vector. Site-directed mutagenesis was used to introduce a silent mutation and eliminate an internal BamHI site generating pTOPO-PTP-ΔBamHI. The Myc tag was replaced with PTP-ΔBamHI to generate pLew100-PTP^Puro^. The full-length open reading frames of wild-type *POLIC* and with the point mutations were cloned into pLew100-PTP^Puro^ (pLewIC-WT-PTP^Puro^). Gibson assembly was used to generate truncated versions of *POLIC* that retained only the C-terminal POLA domain or the N-terminal uncharacterized region (UCR). Briefly, PCR fragments containing the predicted mitochondrial targeting sequence (MTS) (nucleotides 1-174), the POLA fragment (nucleotides 3793-4947), and the UCR fragment (nucleotides 1-3792) were generated. Gibson Assembly (NEB) was then performed using HindIII/XbaI digested pLew100-PTP^PURO^ with the MTS and POLA PCR fragments or with the UCR fragment. All DNA constructs and point mutations were verified by direct sequencing.

##### Trypanosome cell culture and transfection

Procyclic form *T. brucei* strain 29-13 (Wirtz, Leal, Ochatt, Cross, 1999) was cultured at 27°C in SDM-79 medium supplemented with FBS (15%), G418 (15 µg/mL) and hygromycin (50 µg/mL). To generate the parental RNAi cell line IC-UTR, 15 µg of pStLIC3’UTR was transfected by electroporation into 29-13 cells, selected with 2.5 µg/mL phleomycin and cloned by limiting dilution as previously described (Chandler et al 2008). Four individual clones were analyzed and clone P6E2 was chosen based on growth rate, *POLIC* knockdown efficiency, and kDNA defects. All *POLIC* variant expression constructs (15 µg) were transfected by nucleofection using the Amaxa Nucleofection Parasite Kit (Lonza) into 29-13 cells to generate inducible overexpression POLIC-PTP variant cell lines (OE) and into IC-UTR RNAi cell line to generate complementation cell lines (RNAi+OE). Following selection with 1 µg/mL puromycin, cell lines were cloned by limiting dilution. A single clone of each cell line was selected based on variant expression for OE and confirmed *POLIC* knockdown and variant expression for RNAi+OE cell lines. All cell lines were maintained between 1×10^5^ – 1×10^7^, and were not continuously cultured for more than three weeks. Cell density was determined using a Beckman Coulter Z2 particle counter. RNAi and/or overexpression were induced by the addition of tetracycline (1 µg/mL). Cultures were supplemented with 0.5 µg/mL Tet on non-dilution days to maintain POLIC variant expression and/or RNAi (Rusconi, Durand-Dubief, Bastin, 2005).

##### RNA isolation and RT-qPCR analysis

Total RNA was isolated from 5×10^7^ cells with Trizol (Thermo Fisher) using standard procedures. RNA concentration was determined by Nanodrop and 100 ng of total RNA was reverse transcribed to cDNA with random primers and RNase inhibitor (Applied Biosystems). qPCR was performed using QuantiNova SYBR Green PCR (Qiagen) with 1 µg of cDNA and 0.2 µM of primer per reaction (qPCR primers can be found in Table S1) with a Stratagene MxPro 3000x thermocycler. All reactions were performed in triplicate and qPCR results were determined from three biological replicates. Gene knockdown was normalized using *TERT* (Brenndörfer and Boshart, 2010).

##### SDS-PAGE and Western blot analysis

Cells were harvested and washed once with PBS supplemented with protease inhibitor cocktail. Samples were fractionated by SDS-PAGE and transferred overnight onto a PVDF membrane in 1X transfer buffer containing 1% methanol. Membranes were blocked in Tris-buffered saline (TBS) + 5% non-fat dry milk for at least one hour. Peroxidase-Anti-Peroxidase soluble complex (PAP) reagent (1:2000, 30 min) (Sigma, P1291) was used for PTP tag detection. For subsequent detections where stripping was required because proteins of interest were similarly sized to the loading control, membranes were stripped with 0.1 M glycine (pH 2.5, 15 min, 37 °C), washed in TBST (0.1 % Tween-20), blocked and re-probed with specific *Crithidia fasciculata* anti-Hsp70 (1:5000, 1 h) (Johnson and Englund, 1998) and secondary chicken anti-rabbit-HRP (1:2000, 1 h) (Santa Cruz, sc-516087). Signal was detected with Clarity™ ECL Blotting Substrate (NEB) and on a GE Imagequant LAS 4000 mini. Quantification of Western blot intensities was performed using ImageJ software (http://imagej.nih.gov/ij/).

##### DNA isolation and Southern blot analysis

Total DNA was isolated from 1×10^8^ cells using Puregene Core Kit A (Qiagen). Free minicircles and total kDNA content were analyzed as previously described (Bruhn, Mozeleski, Falkin, Klingbeil, 2010). Quantification was performed using a Typhoon 9210 Molecular dynamics Phosphoimager (GE Healthcare) with background subtraction and normalized against the tubulin signal using ImageQuant 5.2 software.

##### Immunoprecipitation and DNA polymerase activity assay

Immunoprecipitation of PTP-tagged POLIC variants was conducted according to (Brandenburg, Schimanski, Nogoceke, Nguyen, Padovan, Chait, Cross, Günzl, 2007) with modifications. Briefly, 3 × 10^8^ mid-log phase cells were harvested (9 × 10^8^ IC-UCR cells were used due to low expression), washed with cold PBS and lysed in 970 μL of 150 mM KCl, 20mM Tris-HCl (pH 8), 3 mM MgCl_2_, 0.5% NP-40, 0.5 mM DTT supplemented with [1X] cOmplete™ EDTA-free protease inhibitor (Roche). Cell lysate was bound to IgG-Sepharose beads (Roche) and washed by centrifugation with 100 mM NaCl, 20 mM Tris-HCl (pH 8), 3 mM MgCl_2_, 0.1% Tween 20 buffer five times at 1,000 x *g* for 5 minutes each. All steps were performed at 4°C. Samples bound to beads were assayed for DNA polymerase activity. Reactions contained 50 mM Tris-HCl (pH 8), 5 mM MgCl_2_, 1 mM DTT, 0.1 mg/mL BSA, 20 μg/mL activated calf thymus DNA as template, 10 μM dTTP, dGTP, and dCTP, 5 μL [α-32P] dATP (Perkin Elmer) mixed with 10 μM cold dATP, either the specific units of Klenow fragment (NEB) or the specified volumes of immunoprecipitated POLIC variants. Reactions were incubated at 37°C for 30 minutes, stopped by adding 5 μL of 500 mM EDTA, and spotted onto Whatman DE81 disks, dried, and then washed four times with 0.5 M NaPO_4_ (15 min each). Disks were then washed with 80% EtOH for 15 min and dried before analyzing with an LS2 Beckman Coulter scintillation counter.

##### Immunofluorescence microscopy

Procyclic cells were pelleted at 975 × *g*, washed and resuspended at approximately 5×0^6^ cells/mL in PBS, then adhered to poly-L-lysine-coated slides (5 min). Cells were then fixed in 4% paraformaldehyde (5 min) and washed three times (5 min) in PBS/0.1 M glycine (pH 7.4). Cells were permeabilized with 0.1% Triton X-100 (5 min) and washed in PBS 3 times (5 min). PTP-tagged proteins were detected by incubating with anti-protein A serum (1:3000, 1 h) (Sigma, P3775) followed by Alexa Fluor® 594 goat anti-rabbit (1:250, 1 h) (Invitrogen, A-11012). Detection of basal bodies (YL1/2), TAC102 and DNA (DAPI) was previously described (Concepción-Acevedo, Miller, Boucher, Klingbeil, 2018). Slides were then washed 3 times (5 min) in PBS prior to mounting in Vectashield (Vector Laboratories). Images were captured at the same exposure time for each channel within the same experiment using a Nikon Eclipse E600 microscope equipped with a cooled CCD Spot-RT digital camera (Diagnostic Instruments) and a 100X Plan Fluor 1.30 (oil) objective. Quantification for kDNA morphology was performed as described previously (Bruhn 2010, Chandler 2008). Asymmetric networks were classified as one cell with two detectable kDNA networks and “other” was classified as a cell with a karyotype that did not match either a normal karyotype (1N1K, 1N2K, 2N2K) or aberrant kDNA morphology (small, none, ancillary, or asymmetric).

## SUPPLEMENTARY DATA

Supplementary Data are available at JSC online.

## ACKNOWLEDGMENTS

We thank Amy Springer for critical review of this manuscript, Paul Englund for the mtHsp70 antibody and Torsten Ochsenreiter for the TAC102 antibody.

---

**Table S1:**
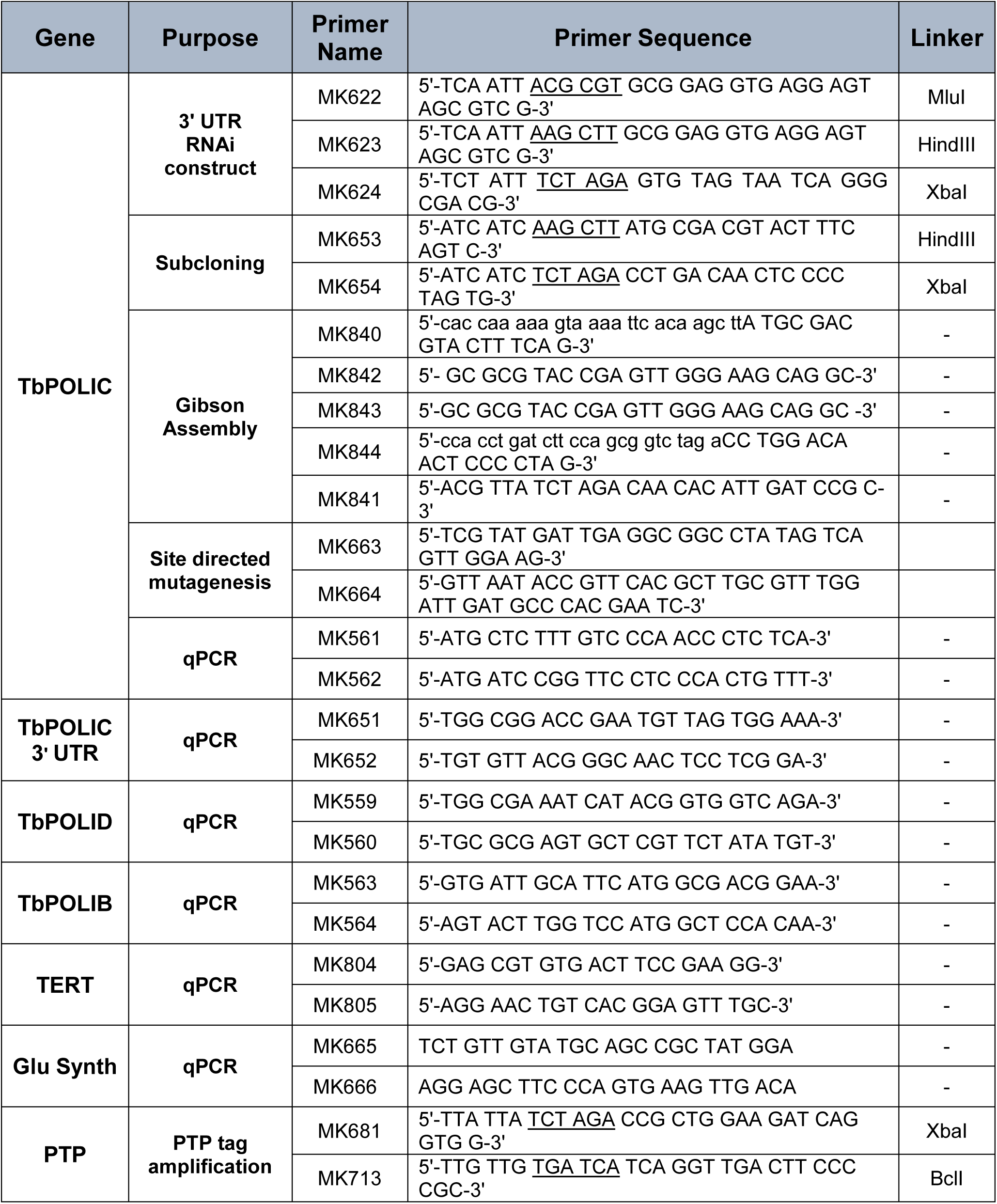
Primers used in this study. Underlined regions correspond to restriction enzyme sites. Lowercase letters correspond to plasmid backbone sequence.

## FIGURE LEGENDS

**Fig S1. TAC102 colocalizes with ancillary kDNA. (A)** Representative images of TAC102 and ancillary kDNA co-localization in the IC-UTR, IC-DEAD RNAi + OE, IC-POLA RNAi + OE, and IC-UCR RNAi + OE cell lines. DAPI-staining (blue); anti-TAC102 (red); YL1/2 (green). The posterior end of the cell is the rounded end and the expected position of normal kDNA. White arrows indicate ancillary kDNA and purple arrowheads show TAC102 colocalized with ancillary kDNA. Scale bar, 5 μm. **(B)** Western blot detection of TAC102 and HSP70 protein levels following 10 days of induction for each of the listed cell lines. 1×10^6^ cell equivalents was loaded into each well except for C where 5×10^6^ cell equivalents was loaded. +C, single expresser control cell line.

**Fig S2. Rescue with ectopically expressed IC-WT in the IC UTR RNAi cell line. (A)** Growth curves of uninduced (white squares), tetracycline-induced (blue diamonds) complementation with IC-WT, and induced parental IC-UTR RNAi (gray squares) procyclic cells over 10 days. Error bars represent the standard deviation from the mean of three biological replicates. **(B)** RT-qPCR analysis of uninduced (black) and induced (blue/white hashed) cells following 48 hours of tetracycline induction. Data are normalized to TERT and error bars represent the standard deviation from the mean of three biological replicates. IC UTR, POLIC 3’ untranslated region; IC ORF, POLIC open reading frame; IB ORF, POLIB open reading frame; ID ORF, POLID open reading frame. **(C)** Western blot detection of the PTP tag and HSP70 over 10 days of induced complementation. At each time point, 1×106 cell equivalents was loaded into each well except for +C where 5×10^6^ cell equivalents was loaded. +C, single expresser control cell line. **(D)** Quantification of kDNA phenotypes over 10 days. Over 300 cells were scored at each time point and error bars represent standard deviation from the mean of three biological replicates. Blue bar, IC-WT complementation; gray bar, POLIC RNAi. Specific kDNA phenotype information can be found in Figure 1E. **(E)** Representative Southern blot showing the effect on free minicircle population. k, kDNA network; N/G, nicked/gapped; L, linearized; CC, covalently closed; T, tubulin. **(F)** Quantification of PTP foci formation at each cell cycle stage. Percentages refer to the fraction of total cells in the population. Green/white hashed, foci positive; gray, foci negative.

**Fig S3. Overexpression of ectopically expressed IC-WT in the absence of *POLIC* RNAi. (A)** Growth curves of uninduced (open circles) and tetracycline-induced (gray circles) procyclic cells over 10 days. Error bars represent the standard deviation from the mean of three biological replicates. **(B)** Western blot detection of the PTP tag and HSP70 protein levels following 10 days of induced overexpression. At each time point, 1×10^6^ cell equivalents was loaded into each well except for C where 5×10^6^ cell equivalents was loaded. +C, single expresser control cell line. **(C)** Representative images of IC-WT-PTP throughout the cell cycle. Scale bar, 5 μm. **(D)** Quantification of PTP foci formation at each cell cycle stage. Percentages refer to the fraction of total cells in the population. Green/white hashed, foci positive; gray, foci negative. **(E)** Representative Southern blot showing the effect on free minicircle population. k, kDNA network; N/G, nicked/gapped; L, linearized; CC, covalently closed; T, tubulin.

**Fig S4. IC-WT displays nucleotide incorporation activity. (A)** Representative immunoblot analyses of 5 µL eluate from POLIC variant immunoprecipitation from three separate experiments. IP products were run on an 8% SDS-PAGE gel, blotted, and detected with PAP reagent (Sigma). 3X value indicates the number of total cells that were isolated for IP. Protein marker masses are indicated in kilodaltons (kDa). **(B)** Representative assay for DNA incorporation activity of Klenow fragment (Invitrogen) and POLIC variants *in vitro* as measured by counts per minute (CPM) in a scintillation counter from three separate experiments. K, Klenow, numbers underneath indicate enzyme units (U) input; WT, IC-WT; DEAD, IC-DEAD; POLA, IC-POLA, UCR, IC-UCR; number indicates µL of eluate input; 29-13, 29-13 cell lysate; Beads, beads only; Buffer, buffer only. Error bars represent standard deviation from two technical replicates.

**Fig S5. Characterization of IC-POLA and IC-UCR overexpression cell lines. (A)** Western blot detection of the PTP tag and HSP70 or alpha tubulin protein levels following 10 days of induction in the OE IC-POLA cell line. **(B)** Quantification of kDNA morphology over the course of the induction in the OE IC-POLA cell line. Over 300 cells were scored at each time point and error bars represent standard deviation from the mean of three biological replicates. Gray, normal kDNA; yellow, small kDNA; red, no kDNA; purple, ancillary kDNA; orange, asymmetric size; green, other. **(C)** Representative Southern blot showing the effect on free minicircle population in the OE IC-POLA cell line. k, kDNA network; L, linearized; N/G, nicked/gapped; CC, covalently closed; T, tubulin. **(D-F)** Same as A-C but for the OE IC-UCR cell line.

